# A comment on *“Species are not most abundant in the centre of their geographic range or climatic niche”*

**DOI:** 10.1101/266510

**Authors:** Jorge Soberón, A. Townsend Peterson, Luis Osorio-Olvera

## Abstract

A study recently published argued against a relationship between population density and position in geographic and environmental spaces. We found a number of methodological problems underlying the analysis. We discuss the main issues and conclude that these problems hinder a robust conclusion about the original question.

## Introduction

The question of whether population density is related to position in geographic (Sagarin 2002) or ecological niche space (Yañez-Arenas et al. 2012, Martínez-Meyer et al. 2013) is important and still unresolved. In a recent paper in *Ecology Letters*, Dallas et al. (2017) examined the problem using a large dataset of 118,000 sampled populations of >1400 species birds, mammals, and trees. Dallas et al. (2017) failed to detect consistent and significant correlations between population density and distance to the centroids of species’ distributions in geographic or environmental spaces, and concluded against the generality of such distance-density relationships. However, the authors’ failure to detect significant relationships may result from methodological artifacts, rather than to non-existence of such relationships. We focus on five problems inherent in their analysis.

## Results

1) The largest dataset analyzed by Dallas et al. (2017) was eBird observations (Sullivan et al. 2009), which are collected without any sampling protocol or plan (there are alternative and better databases, like the Breeding Bird Survey). eBird has biases frequent among observational data, like more observers near cities, and more reporting where a species is rare. Therefore, confounding effects between effort and observer bias may be present, at least for the birds.

2) Dallas et al. (2017) caution about maximum abundances falling at the periphery of sampled ranges for two of the datasets that they analyzed, but we still worry that true niche centroids will not be represented appropriately. Dallas et al. (2017) largely disregarded parts of species’ distributions falling outside the regions for which they had abundance data available. We illustrate this point using the rodent *Dipodomys merriami* Dallas et al. (2017). Figure 1 shows the spatial minimum convex hull (CH) for occurrences in the United States (region in gold on map below). This is considerably less extensive than the range outline for this species from IUCN (Patterson et al. 2003). The geographic centroids based on the two range outlines are markedly distinct.

**Figure 1.**
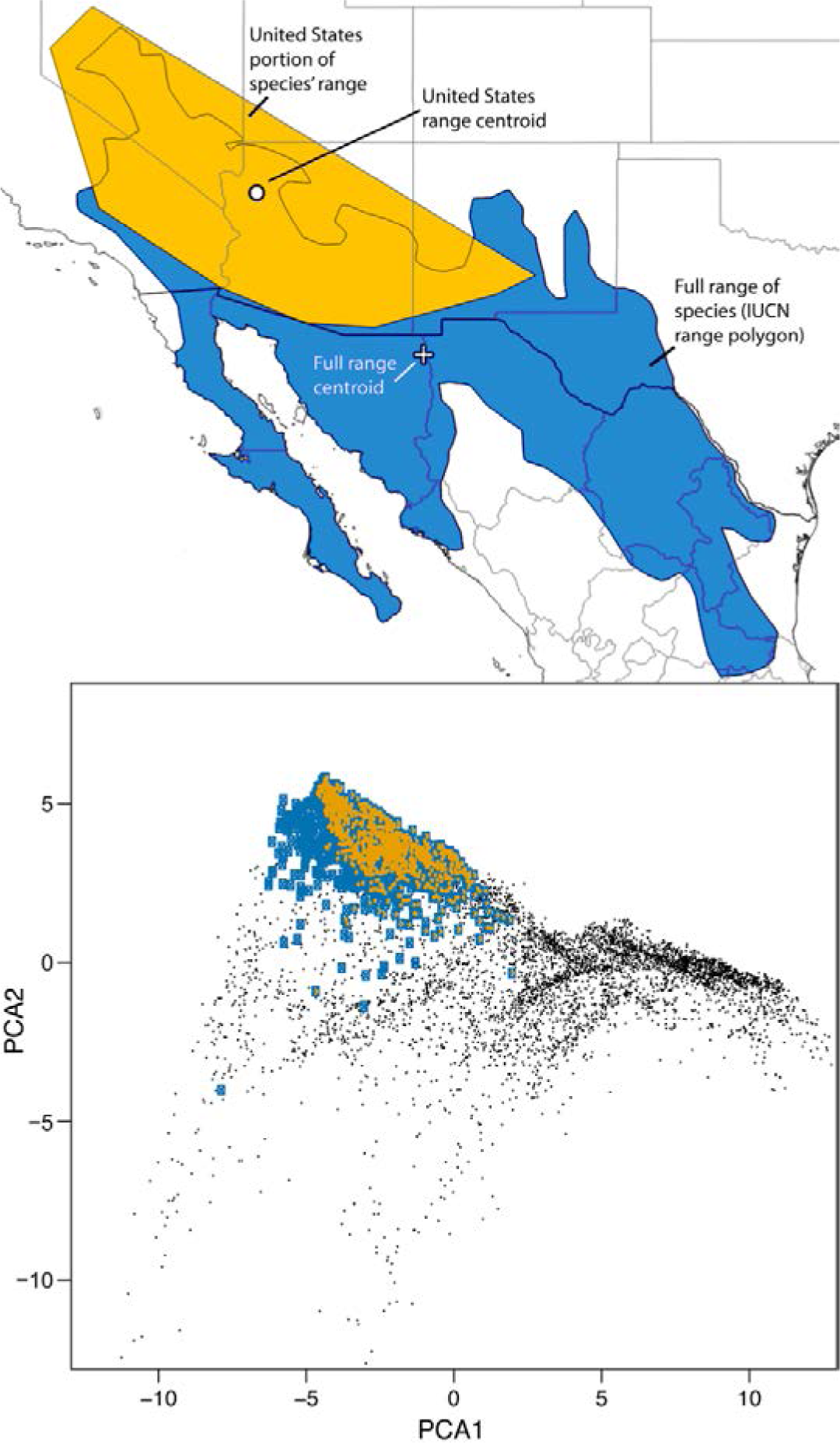
Geographic and environmental spaces for *Dipodomys merriami*. Top: extent of occurrence polygon for the distribution of *D. merriami* in the United States (gold, centroid shown by black and white circle), representing the range area analyzed by Dallas et al. (2017), and the full range of the species (blue, centroid show as a cross; polygon from IUCN). Note that the true range centroid falls outside of the convex hull analyzed by Dallas et al. (2017). Bottom: 1799 data points from GBIF (see text) in a space of the first two climatic principal components used by Dallas. et al. (2017; see text). Points in gold are the reduced portion (United States) of the species range analyzed by Dallas et al. (2017); points in in blue cover the entire range of the species.

A similar problem exists in environmental space. We downloaded the 2-dimensional principal components (PC) used by Dallas et al. (2017). For 1799 localities (debugged and thinned to 0.1°, out of 40,000 available via GBIF), we extracted the PC values for each of the points. Figure 1 shows that the range of environmental space in the full distributional area of *Dipodomys merriami* extends into environmental space not represented in the CH used by Dallas et al. (2017).

3) Dallas et al. (2017) used CHs to characterize ecological niches of species. CHs are sensitive to outliers (Syväranta et al. 2013), and their centroids may be quite distinct from those obtained using robust estimators (Van Aelst and Rousseeuw 2009). In Figure 2, based on the *D. merriami* example, the CH and a minimum volume ellipsoid (MVE) centroids around the same US data are located in very different positions in niche space.

**Figure 2.**
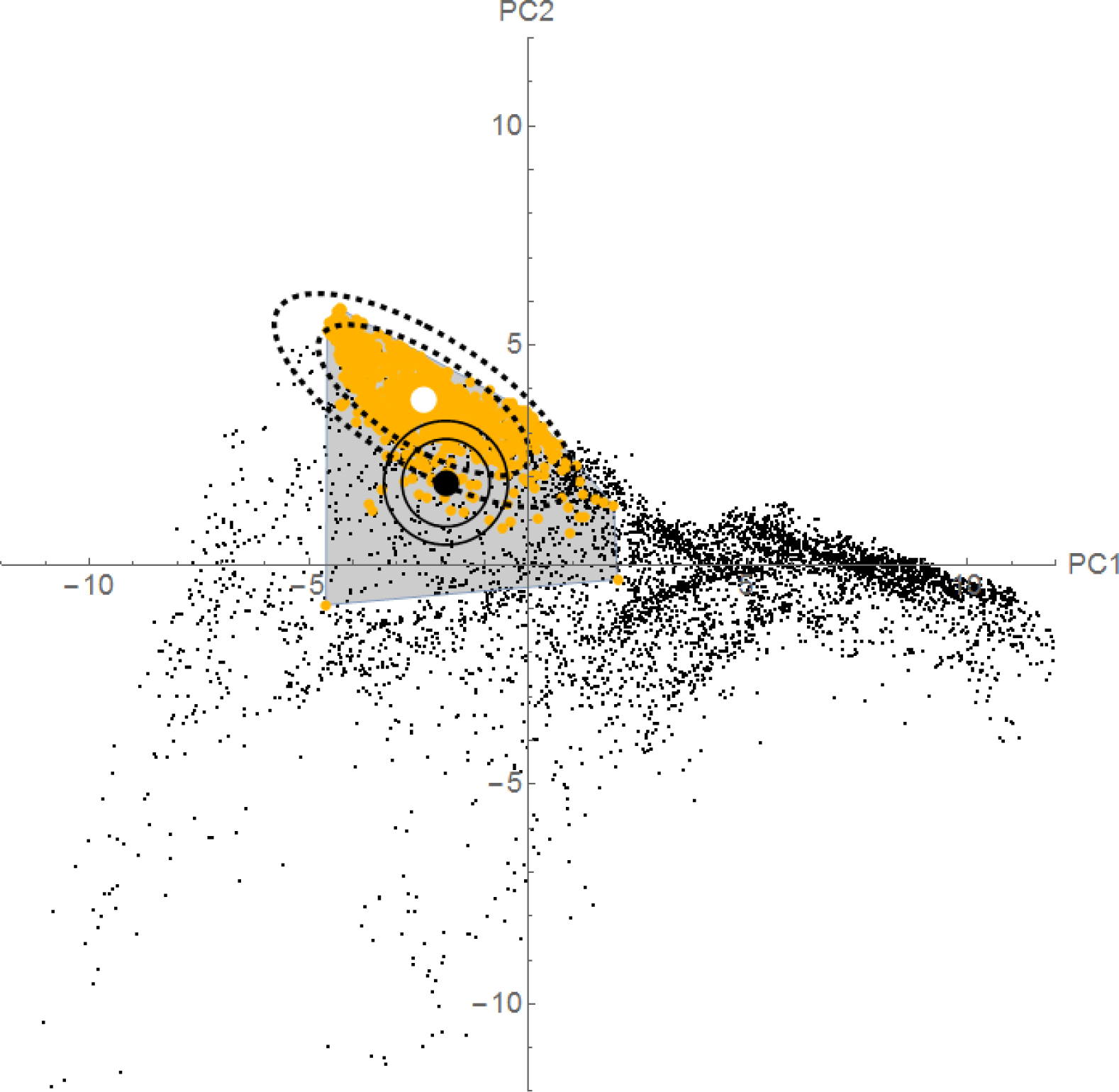
GBIF points for *Dipodomys merriami* in environmental space, showing differences between methods for delimiting niches and calculating niche-centroid distances. The black circle is the centroid of the convex hull (gray-shaded polygon), showing the strong effect of one outlier point. The white circle is the centroid of a 95% minimum volume ellipsoid that is able to ignore the outlier. Circles are Euclidean distances of radii 1 and 2, for the convex hull centroid; the dashed ellipsoids are the equivalent distances (Mahalanobis distances) taking into account the covariance shown by the points in gold (see text). Note the striking differences between the two methodologies in both shape of the niche estimated and the distances that result; in particular, note that the centroid estimated via convex hulls falls at the periphery of the cloud of points for the species’ occurrence.

4) Dallas et al. (2017) use Euclidean distances as measures of distance to niche centroids, which trace equidistant circles around the centroid. A Mahalanobis distance, estimated using the covariance matrix of the observations of the species in question, would be preferable. Figure 2, shows the US distribution of *D. merriami* occurrences, with centroids and outlines of the CH and MVE. Ignoring the covariance in the realized niche of the species contributes extra bias to characterizations of distances in niche space. Dallas et al. (pers. comm.) explored Mahalanobis distances, without finding major differences in the results, an observation deserving further exploration.

**5)** The data provided in the Supplementary Materials of Dallas et al. (2017) reveal that the population-density data points have coordinates with precision of 100 m or finer. However, the climate data used in the paper have a resolution of ∼0.042°, or squares of ∼4600 m on a side. This means that multiple abundance data points may fall within a single climate pixel, introducing a further problem in the analysis, as shown in lines 365-370 of the code provided by the authors. Correcting this methodological problem leaves 40 instead of 81 species for mammals, 49 instead of 63 for fishes with >10 different abundance/climate points; all of the birds and 165 of the 166 tree species.

## Conclusions

Dallas et al. (2017) provide the largest-scale analysis available to date of relationships between population density and positions in geographic and environmental spaces. Their negative results contrast with previous empirical work (Yañez-Arenas et al. 2012, Martínez-Meyer et al. 2013) and with theoretical arguments supporting such a relationship (Maguire 1973, Osorio-Olvera et al. 2016).

In this communication, we identify a series of methodological problems underlying the results of Dallas et al. (2017). Although some of them may be of minor importance (e.g., Euclidean vs. Mahalanobis distances), others may be more significant. We suggest that this important and interesting problem remains far from settled.

